# *In vivo* PSC differentiation as a platform to identify factors for improving the engraftability of cultured muscle stem cells

**DOI:** 10.1101/2023.12.26.573361

**Authors:** Ning Xie, Kathryn Robinson, Timothy Sundquist, Sunny S. K. Chan

**Author notes:** Correspondence: Sunny S.K. Chan.

## Abstract

Producing an adequate number of muscle stem cells (MuSCs) with robust regenerative potential is essential for the successful cell therapy of muscle-wasting disorders. We have recently developed a method to produce skeletal myogenic cells with exceptional engraftability and expandability through an *in vivo* pluripotent stem cell (PSC) differentiation approach. We have subsequently mapped engraftment and gene expression and found that leukemia inhibitory factor receptor (*Lifr*) expression is positively correlated with engraftability. We therefore investigated the effect of LIF, the endogenous ligand of LIFR, on cultured MuSCs and examined their engraftment potential. We found that LIF-treated MuSCs exhibited elevated expression of PAX7, formed larger colonies from single cells, and favored the retention of PAX7^+^ “reserve cells” upon myogenic differentiation. This suggested that LIF promoted the maintenance of cultured MuSCs at a stem cell stage. Moreover, LIF enhanced the engraftment capability of MuSCs that had been expanded *in vitro* for 12 days by 5-fold and increased the number of MuSCs that repopulated the stem cell pool post-transplantation. These results thereby demonstrated the effectiveness of our *in vivo* PSC differentiation platform to identify positive regulators of the engraftability of cultured MuSCs.

## 1. Introduction

Cell therapy is an attractive therapeutic strategy for chronic diseases due to its promise to replace damaged tissues with new healthy donor cells. For Duchenne muscular dystrophy (DMD) which can be caused by any 1 of the more than 2000 different mutations in the DMD gene, cell therapy offers potential benefits regardless of the exact genetic mutation which can vary from patient to patient (Biressi et al., 2020; Blau and Daley, 2019). However, DMD cell therapy has its own unique challenges. A prominent problem is the difficulty of obtaining an ideal donor cell type that is both expandable to vast amounts for clinical use and engraftable to form new fibers after transplantation (Biressi et al., 2020; Blau and Daley, 2019; Verhaart and Aartsma-Rus, 2019). Muscle stem cells (MuSCs), also known as satellite cells, are an endogenous cell population responsible for maintaining the lifetime integrity of skeletal muscles against wear and tear (Günther et al., 2013; von Maltzahn et al., 2013). MuSCs have tremendous regenerative potential *in vivo*, with a single MuSC capable of regenerating hundreds of fibers (Collins et al., 2005; Sacco et al., 2008). Nevertheless, their clinical applications remain limited due to 2 main reasons. First, MuSCs are scarce and cannot be obtained at a therapeutically meaningful quantity from small muscle biopsies (Roth et al., 2000). An *in vitro* expansion step is inevitable. Second, MuSCs after *in vitro* expansion lose their regenerative potential dramatically, such that transplantation of hundreds of thousands of expanded MuSCs can merely give rise to a few hundred fibers (Montarras et al., 2005; Sacco et al., 2008; Xie et al., 2021). There is an urgent need to produce MuSCs that are both expandable and engraftable.

We have recently developed an *in vivo* pluripotent stem cell (PSC) differentiation method to produce skeletal myogenic cells that are both expandable and engraftable (Chan et al., 2018; Xie et al., 2021; Xie et al., 2023). These skeletal myogenic cells are highly efficient in forming new fibers and reconstituting the MuSC niche upon transplantation, and they can be expanded *in vitro* for over a month while maintaining high engraftability (Chan et al., 2018; Xie et al., 2021; Xie et al., 2023). Moreover, the development of the skeletal myogenic lineage during *in vivo* PSC differentiation closely recapitulates embryonic skeletal myogenesis (Pappas et al., 2022). These observations therefore suggest that *in vivo* PSC differentiation is effective not only as a method to produce engraftable skeletal myogenic cells, but also as a unique platform to study how the engraftability of skeletal myogenic cells is determined. Given the unmet challenge of producing a skeletal myogenic cell type that is both engraftable and expandable, *in vivo* PSC differentiation offers an invaluable approach to discover factors that promote the engraftability of expanded MuSCs.

In the current study, we have used *in vivo* PSC differentiation to identify leukemia inhibitory factor (LIF) as a potential regulator of skeletal myogenic engraftment. We subsequently validated our discovery in showing that LIF substantially improved the engraftment potential of expanded MuSCs that have been cultured for 12 days over 3 passages.

## 2. Materials and Methods

### Animals

All procedures involving animals including animal housing, husbandry, and experiments were reviewed and approved by the University of Minnesota Institutional Animal Care and Use Committee with AAALAC accreditation according to protocols (#2201-39776A). Male and female wildtype BL6 (C57BL/6J, Jackson Laboratory, Bar Harbor, ME) and H2B-GFP mice (B6.Cg-Tg(HIST1H2BB/EGFP)1Pa/J, Jackson Laboratory) at 3-5 months old were used to obtain MuSCs. Transplantation experiments were performed on both male and female 3-5 months old NSG-mdx^4cv^ mice as described previously (Arpke et al., 2013).

### Muscle stem cell isolation

Hindlimb muscles were harvested and chopped into ∼2 mm pieces. The chopped muscle pieces were then incubated in a primary digestion buffer containing Dulbecco’s Minimum Essential Medium/High Glucose (DMEM, HyClone #SH30243.01, Logan, UT), 2 mg/mL Collagenase II (Gibco #17101-015, Gaithersburg, MD), and 1% penicillin/streptomycin (P/S) (Life Technologies #15140-122, Grand Island, NY) on a shaker at 250 rpm, 37 °C for 1 hour. Primary digestion was then halted by the addition of rinsing buffer consisting of Ham’s/ F-10 medium (Caisson Labs #HFL01, Smithfield, UT), 10% horse serum (HyClone #SH30074.03), 1% HEPES buffer solution (Caisson Labs #HOL06), and 1% P/S and then centrifuged at 500 g for 10 minutes at 4°C. The tissues were then subjected to further enzymatic digestion consisting of rinsing buffer supplemented with 0.1 mg/mL Collagenase II and 0.5 mg/mL Dispase (Gibco #17105-041). This secondary digestion process continued for 30 minutes on a shaker at 250 rpm, 37 °C. The digested tissues were then repeatedly drawn and released into a 10 mL syringe with a 16-gauge needle (4 times) followed by an 18-gauge needle (4 times) to facilitate additional dissociation. The resultant cellular suspension was filtered through a 100 μm cell strainer, spun down, resuspended in rinsing buffer, filtered through a 40 μm cell strainer, and then spun down again. Isolation of muscle stem cells from transplanted tibialis anterior muscles (see below) was performed similarly, except without passing through the 16-gauge needle nor the 100 μm cell strainer.

### Fluorescence-activated cell sorting (FACS)

Isolated cells were incubated on ice for one hour with fluorophore-conjugated antibodies for FACS (fluorescence-activated cell sorting). After one hour, the cells were washed twice and resuspended in FACS buffer (PBS (HyClone #SH30256.01), 0.2% fetal bovine serum (FBS), and 1 μg/mL propidium iodide (PI, Sigma-Aldrich #P4170, St Louis, MO). The addition of PI into FACS Buffer served as a live/dead cell indicator, and only viable cells (PI^−^) were quantified. Cells were sorted into medium and kept on ice until they were cultured as described below. The antibodies utilized for sorting muscle stem cells (each at concentration of 0.5 μL per million cells) were PE-Cy7 anti-CD31 (BioLegend #102418, RRID: AB_830757, San Diego, CA), PE-Cy7 anti-CD45 (BioLegend #103114, RRID: AB_312979), APC anti-α7-Integrin (AbLab #67-0010-05, Vancouver, Canada), and PE anti-VCAM-1 (BioLegend #105714; RRID: AB_1134164). MuSCs are defined as CD31^−^ CD45^−^ α7-Integrin^+^ VCAM-1^+^ (α7-Integrin^+^ VCAM^+^). Cell analysis and sorting were executed using BD FACSAriaII instrument (BD Biosciences, San Diego, CA) operated with FACSDiva software (BD Biosciences). A four-way purity precision was employed for bulk sorting, while single-cell precision was implemented when sorting individual cells into 96-well plates for clonal analysis. FACS plots depicting the distribution of cellular populations were generated with FlowJo software (FLOWJO LLC, Ashland, OR).

### In vitro cell expansion and differentiation

FACS-sorted α7-Integrin^+^ VCAM^+^ cells were plated in 0.1% gelatin-coated wells and cultured in myogenic expansion medium (Ham’s/F-10, 20% FBS (Sigma-Aldrich #F0926), 10 ng/mL basic FGF (R&D Systems #233-FB/CF, Minneapolis, MN), 1% P/S, 2 mM Glutamax (Life Technologies #35050-061, Paisley, PA), and 0.1 mM β-mercaptoethanol) with or without LIF (1000 units/mL, Sigma-Aldrich #ESG1107). Cells were passaged once they reached ∼60% confluency. For differentiation experiments, cells at 70-80% confluency were switched to myogenic differentiation medium (DMEM, 4% horse serum, 2 mM Glutamax, 1 mM sodium pyruvate (Life Technologies #11360070), 1 μg/mL insulin (GeminiBio #800122, West Sacramento, CA), 1 μM dexamethasone (Hello Bio #HB2521, Princeton, NJ), and 1% P/S) for 3 days.

### Clonal analysis

One α7-Integrin^+^ VCAM^+^ cell per well was seeded via FACS into 0.1% gelatin-coated 96-well plates in myogenic clonal medium containing DMEM/F12 (Cellgro #15-090-CV, Manassas, VA), 20% FBS, 10% horse serum, 10 ng/mL basic FGF, 1% P/S, 2 mM Glutamax, and 0.5% chick embryo extract (US Biological #C3999, Salem, MA) with or without LIF (1000 units/mL). Cells were incubated for 7 days undisturbed before analysis.

### Cell transplantation

Two days prior to cell transplantation, the hindlimbs of recipient NSG-mdx^4cv^ mice were exposed to 1200 cGy X-Ray irradiation. One day prior to transplantation, the recipients’ irradiated tibialis anterior (TA) muscles were injected with 15 μm of cardiotoxin (10 μM, Latoxan #L8102, France) to promote engraftment. For each recipient mouse, 40,000 donor cells were resuspended in PBS (total 10 μL) and injected into the TA muscle using a Hamilton syringe (Hamilton, Reno, NV). Recipients were anesthetized with ketamine (150 mg/kg, Dechra Veterinary Products, NDC: 59399-114-10, Overland Park, KS) and xylazine (10 mg/kg, Bimeda, NDC: 59399-111-50, Cambridge, Canada) intraperitoneally prior to each procedure. Transplanted TA muscles were harvested 4 weeks later for further analysis.

### Sectioning of transplanted TA muscles

Harvested TA muscles were embedded in optimal cutting temperature (OCT) solution (Scigen #4586, Gardena, CA). The cryomold containing the specimens were snap-frozen on 2-methylbutane (Sigma-Aldrich #320404) pre-cooled with liquid nitrogen and stored at −80 °C. Cryosections of 10 μm were cut on a Leica CM3050 S cryostat (Leica Microsystems, Buffalo Grove, IL) and collected onto glass slides for immunostaining.

### Immunostaining on cultured cells and muscle sections

For cultured cells, cells were fixed with 4% paraformaldehyde (Sigma-Aldrich #P6148) for 1 hour. Cell membranes were permeabilized with 0.3% Triton X-100 (Sigma-Aldrich #X100) followed by blocking with 3% BSA (Thermo Fisher Scientific #BP1605, Waltham, MA). Cells were incubated overnight at 4 °C with primary antibodies in 3% BSA. The next day, cells were incubated for 1 hour at room temperature with secondary antibodies. Nuclei were stained with 4’, 6-diamidino-2-phenylindole (DAPI) (Life Technologies #D3571) for 10 min. For sections, samples were mounted with Immu-Mount (Thermo Fisher Scientific #9990402). Primary antibodies used were mouse anti-PAX7 (1:10, Developmental Studies Hybridoma Bank (DSHB) #PAX7, RRID: AB_395942, Iowa City, IA), mouse anti-myosin heavy chain (MHC) (1:20, DSHB #MF-20, RRID: AB_427788), mouse anti-MYOD1 (1:100, BD Pharmingen #554130, RRID: AB_395255, Franklin Lakes, NJ), rabbit anti-DYSTROPHIN (1:250, Abcam #ab15277; RRID: AB_301813, Cambridge, United Kingdom), and mouse anti-laminin (1:500, Sigma-Aldrich #L8271; RRID: AB_477162). Secondary antibodies (each 1:1000) used were: goat anti-mouse IgG1 Alexa Fluor 555 (Life Technologies, #A21127, RRID: AB_2535769), goat anti-mouse IgG2b Alexa Fluor 647 (Life Technologies, #A21242, RRID: AB_ 2535811), goat anti-rabbit Alexa Fluor 555 (Life Technologies, #121429, RRID: AB_2535850), goat anti-mouse Alexa Fluor 647 (Life Technologies, #A21235, RRID: AB_2535804), and goat-anti mouse IgG2b Alexa Fluor 488 (Life Technologies, #121141, RRID: AB_ 2535778).

### Imaging and analysis

Fluorescent imagery was acquired utilizing a Zeiss AxioObserver Z1 inverted microscope paired with an AxioCamMR3 camera (Jena, Germany). Subsequent image processing and quantification procedures were executed with Fiji/ImageJ software (U.S. National Institute of Health, Bethesda, MD). Whole TA images were captured in tile mode and stitched together using ZEN software (Zeiss). Fiber engraftment is defined as the total cross-sectional area of DYSTROPHIN^+^ fibers over the total cross-sectional area of the whole TA, measured via Muscle2View, a CellProfiler pipeline (Sanz et al., 2019), with adjusted parameters.

### Quantitative RT-PCR

Cells at 70-80% confluency were harvested for gene expression analysis. Total RNA was extracted using Quick-RNA Miniprep Kit (Zymo Research #R1055, Orange, CA). Genomic DNA removal and reverse transcription (RT) were performed using Verso cDNA Synthesis Kit (Thermo Fisher Scientific #AB1453A). Quantitative reverse transcription polymerase chain reaction (qRT-PCR) was performed in triplicates using TB Premix Ex Taq II (Takara Bio, #RR390A, Japan) in a QuantStudio 6 Flex Real-Time PCR System using QuantStudio Real-Time PCR Software (both Applied Biosystems, Waltham, MA). TaqMan probes used were: *Pax7*: Mm00834079_m1, *Myod1*: Mm00440387_m1, *Myog*: Mm00446194_m1, and *Gapdh*: Mm99999915_g1 (all Thermo Fisher Scientific). The ΔΔCt method was utilized to calculate gene expression relative to that of the housekeeping gene *Gapdh* in control samples.

### Statistical analysis

The RNA-seq dataset in Figure 1 was assessed from GEO: GSE182508. Data are presented as mean ± SEM. Differences between two groups were assessed using Student’s t test. Differences among three or more groups were assessed by ANOVA with Tukey’s post-hoc tests. p values < 0.05 were considered significant (*: p < 0.05, **: p < 0.01, and ***: p < 0.001). GraphPad Prism (GraphPad Software, La Jolla, CA) was used to perform statistical analyses.

**Figure 1.**
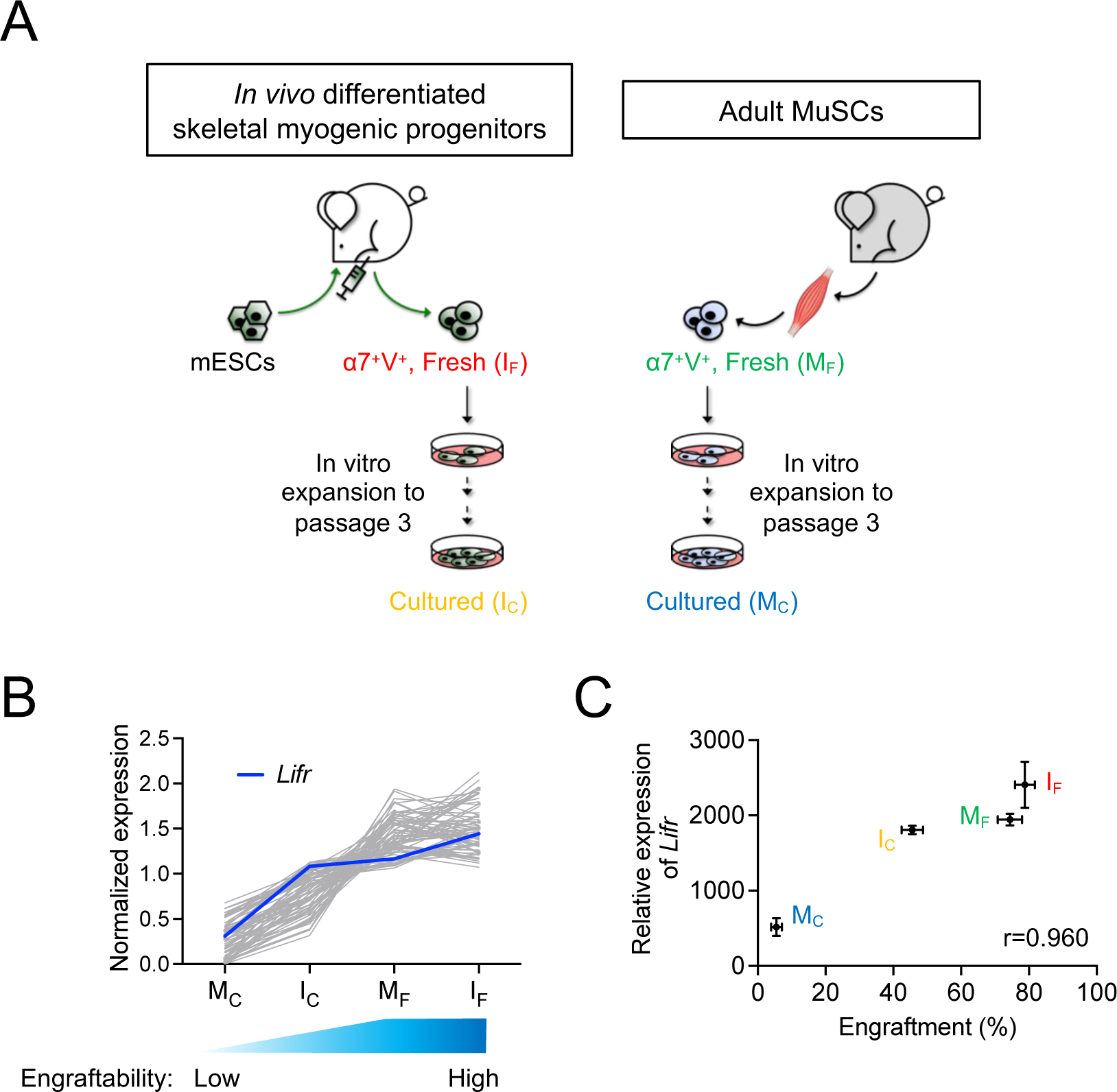
*Lifr* expression was positively correlated with the engraftability of skeletal myogenic cells. (**A**) Schematic of the 4 skeletal myogenic populations (α7-Integrin^+^ VCAM^+^, or α7^+^V^+^) used for RNA-seq analysis: freshly isolated (I_F_) and cultured (I_C_) *in vivo* differentiated skeletal myogenic progenitors and freshly isolated (M_F_) and cultured (M_C_) adult MuSCs (2 biological replicates from each group) (Xie et al., 2021). (**B**) Expression of genes with a Pearson correlation coefficient of > 0.9 (gray lines). Expression of *Lifr* is shown in blue. See text for details. (**C**) Pearson correlation coefficient (r) between engraftment (%) and *Lifr* gene expression (MRN-normalized read counts) from the 4 cell populations.

## 3. Results

### 3.1 Lifr expression was positively correlated with the engraftability of skeletal myogenic cells

We previously reported that *in vivo* PSC differentiation from both mouse and human PSCs produced skeletal myogenic progenitors that were both engraftable and expandable (Chan et al., 2018; Pappas et al., 2022; Xie et al., 2021; Xie et al., 2023). These *in vivo* differentiated skeletal myogenic progenitors had excellent muscle regeneration potency upon transplantation and could produce new fibers to a similar extent as adult MuSCs (Chan et al., 2018; Xie et al., 2021). Remarkably, *in vivo* differentiated skeletal myogenic progenitors could be cultured and expanded *in vitro* over several passages while still retaining their outstanding engraftability (Xie et al., 2021; Xie et al., 2023). This contrasts with adult MuSCs whose engraftment potential abruptly diminished upon a few days in culture (Montarras et al., 2005; Sacco et al., 2008; Xie et al., 2021). We speculated that factors that regulate engraftment might have their expression level correlating to engraftability. In other words, cell populations with better engraftment would express pro-engraftment factors at higher levels, and vice versa. Therefore, in a previous study, we performed an RNA-seq analysis together with a transplantation assay to evaluate the relationship between gene expression and fiber engraftment on 4 skeletal myogenic populations with various degrees of engraftability: fresh MuSCs (M_F_), cultured MuSCs (M_C_), fresh *in vivo* differentiated skeletal myogenic progenitors (I_F_), and cultured *in vivo* differentiated skeletal myogenic progenitors (I_C_) (Figure 1A) (Xie et al., 2021). We first calculated the Pearson correlation coefficient between gene expression and engraftment and identified genes with a positive correlation of r > 0.9 (Figure 1B). We were particularly interested in factors involved in signaling transduction pathways because their activities would be more readily modulated by commercially available agonists and inhibitors. Using these criteria, we identified *Lifr* (LIFR): highly expressed in I_F_ and M_F_ (both highly engraftable), moderately expressed in I_C_ (moderately engraftable), and minimally expressed in M_C_ (least engraftable) (Figure 1B-C).

### 3.2 LIF promoted Pax7 expression in cultured MuSCs

LIFR is a receptor of LIF and the LIF-LIFR pathway has been shown to regulate MuSC biology (Bower et al., 1995; Kami et al., 2000; Spangenburg and Booth, 2002; White et al., 2001). When MuSCs were isolated from the hind limb muscles of adult mice and cultured *in vitro* for 4 days, the skeletal muscle-specific transcription factor MYOD1 emerged with a concurrent downregulation of the MuSC factor PAX7 (Figure 2A-B). This corresponds to the spontaneous differentiation of MuSCs into a PAX7^−^ MYOD1^+^ myoblast stage *in vitro*, where their engraftment potential is severely limited (Olguin and Olwin, 2004; Sacco et al., 2008; Zammit et al., 2004). Interestingly, LIF (1000 units/mL) upregulated PAX7 while producing minimal effect on MYOD1 (Figure 2A-B). These observations were also supported by gene expression analysis in which MuSCs cultured with LIF exhibited elevated levels of *Pax7*, while both *Myod1* and *Myog* expression remained unchanged (Figure 2C).

**Figure 2.**
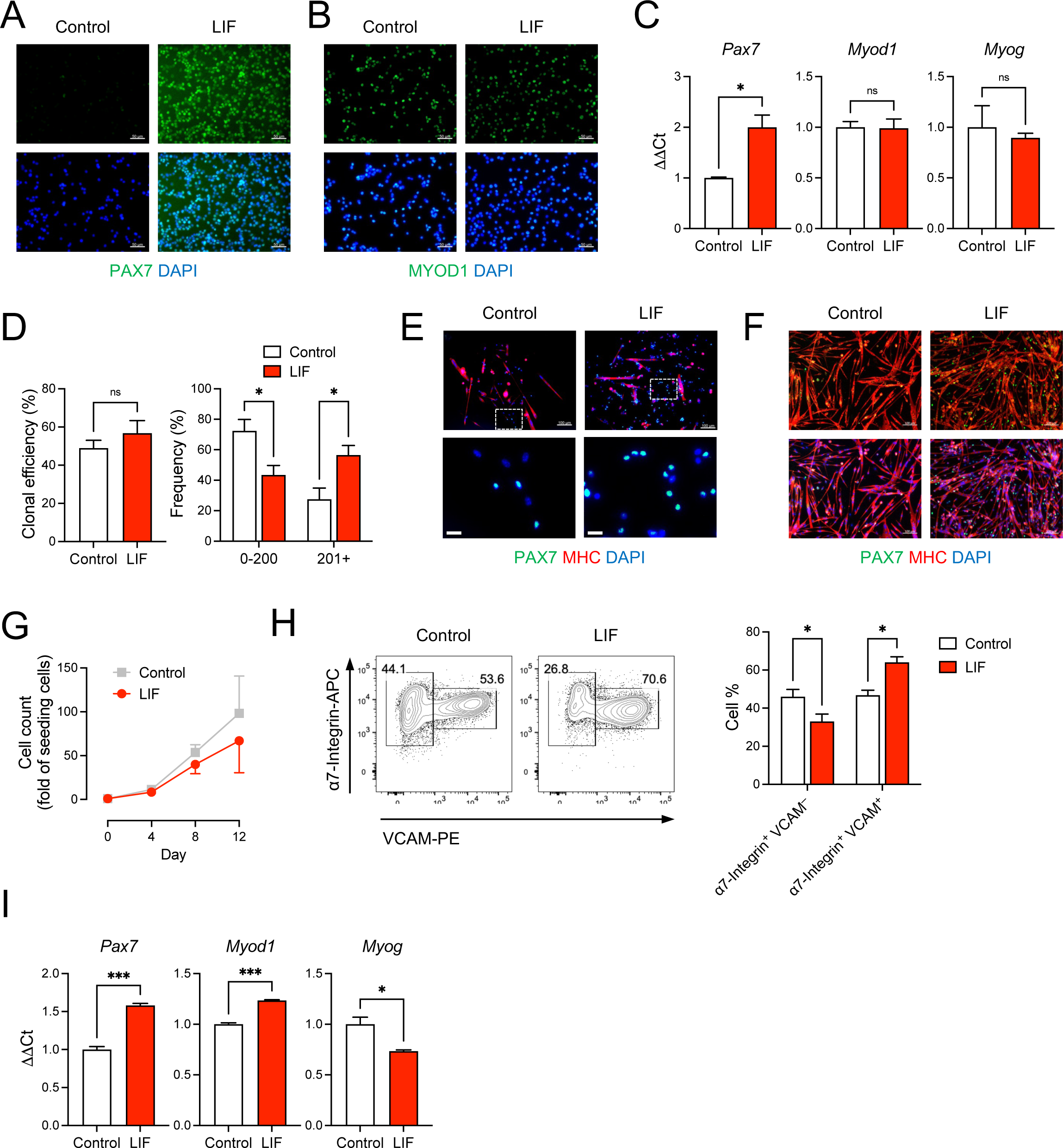
LIF promoted the maintenance of cultured MuSCs at a stem cell stage. (**A,B**) Immunostaining of (**A**) PAX7 and (**B**) MYOD1 in MuSCs cultured *in vitro* with or without LIF for 4 days (scale bar = 50 μm). (**C**) qRT-PCR analysis of cultured MuSCs (mean ± SEM, 3 technical replicates, * p < 0.05). (**D,E**) Clonal analysis of individual MuSCs showing that (**D**) LIF promoted larger colonies (mean ± SEM, 6 biological replicates, * p < 0.05) and (**E**) with more PAX7 expression (top: scale bar = 100 μm). Magnified images were shown at the bottom (bottom: scale bar = 20 μm). (**F**) Upon differentiation, most cultured MuSCs developed into MHC^+^ multi-nucleated myotubes while some became PAX7^+^ (“reserve cells”). LIF promoted the “reserve cell” population (scale bar = 100 μm). (**G**) Growth curves of MuSCs cultured for 12 days over 3 passages with or without LIF. (**H**) FACS analysis of passage 3 MuSCs: (left) typical plots and (right) quantification (mean ± SEM, 3 biological replicates, * p < 0.05). (**I**) qRT-PCR analysis of passage 3 MuSCs (mean ± SEM, 3 technical replicates, * p < 0.05, *** p < 0.001).

We further evaluated how LIF influenced the colony-forming ability of individual MuSCs. Single α7-Integrin^+^ VCAM^+^ MuSCs were FACS-sorted and cultured for 7 days under conditions that promoted both self-renewal and differentiation. Although LIF had minimal impact on the capability of single MuSCs to form myogenic colonies per se, LIF-treated MuSCs tended to develop into larger colonies (Figure 2D-E).

Under pro-differentiation conditions, isolated MuSCs gradually developed into MHC^+^ multi-nucleated myotubes (Figure 2F). Nevertheless, a small population of MuSCs did not differentiate but remained PAX7^+^, known as “reserve cells” (Day et al., 2010; Olguin and Olwin, 2004). Interestingly, we observed an abundance of PAX7^+^ “reserve cells” in LIF-treated MuSC undergoing differentiation (Figure 2F). The higher level of PAX7 in MuSCs treated with LIF under both maintenance and differentiation conditions suggested that LIF might promote cultured MuSCs to remain at the stem cell stage, and thereby making them more likely to be engraftable.

The above experiments were performed on MuSCs cultured for a relatively short period of time (e.g., 4 days) and without passaging. We next evaluated the effect of LIF on MuSCs that have been cultured for 12 days over 3 passages. We did not observe a significant difference in cell growth between control and LIF-treated MuSCs, although the latter tended to proliferate slower (Figure 2G). Interestingly, LIF promoted the α7-Integrin^+^ VCAM^+^ population (markers of MuSCs) in day 12/passage 3 MuSCs at the expense of the α7-Integrin^+^ VCAM^−^ population (markers of myoblasts) (Figure 2H). Moreover, similarly to the effect on the unpassaged MuSCs, LIF upregulated *Pax7* gene expression in passage 3 MuSCs (Figure 2I, left panel). In addition, LIF-treated passage 3 MuSCs exhibited higher levels of *Myod1* with lower levels of *Myog* than untreated controls (Figure 2I, middle and right panels). Altogether, these findings suggested that LIF promoted the maintenance of cultured MuSCs at a stem cell stage.

### 3.3 LIF enhanced the engraftability of cultured MuSCs

The above results encouraged us to evaluate whether LIF could improve the engraftment potential of cultured MuSCs in a transplantation assay (Figure 3A). We first obtained MuSCs from the hind limb muscles of wildtype BL6 mice by FACS using established markers: CD31^−^ CD45^−^ (i.e., non-endothelial and non-hematopoietic, respectively) α7-Integrin^+^ VCAM^+^. We subsequently cultured isolated α7-Integrin^+^ VCAM^+^ cells in myogenic medium with or without LIF (1000 units/mL) for 12 days over 3 passages. Untreated and LIF-treated passage 3 MuSC cultures were then transplanted at 40,000 cells/muscle into the TA muscles of NSG-mdx^4Cv^ mice, a DMD model we have previously used to evaluate cell transplantation success (Arpke et al., 2013; Chan et al., 2018; Xie and Chan, 2023; Xie et al., 2021; Xie et al., 2023). Four weeks later, the transplanted muscles were harvested for fiber engraftment evaluation, as defined by donor-derived DYSTROPHIN^+^ fibers (recipient mice lack dystrophin in their muscles). As expected, untreated cultures engrafted poorly (3.5 ± 0.8%, n = 8). Remarkably, LIF treatment significantly increased the ability of passage 3 MuSCs to develop into DYSTROPHIN^+^ fibers upon transplantation (18.2 ± 3.2%, n = 8, p < 0.001) (Figure 3B-C).

**Figure 3.**
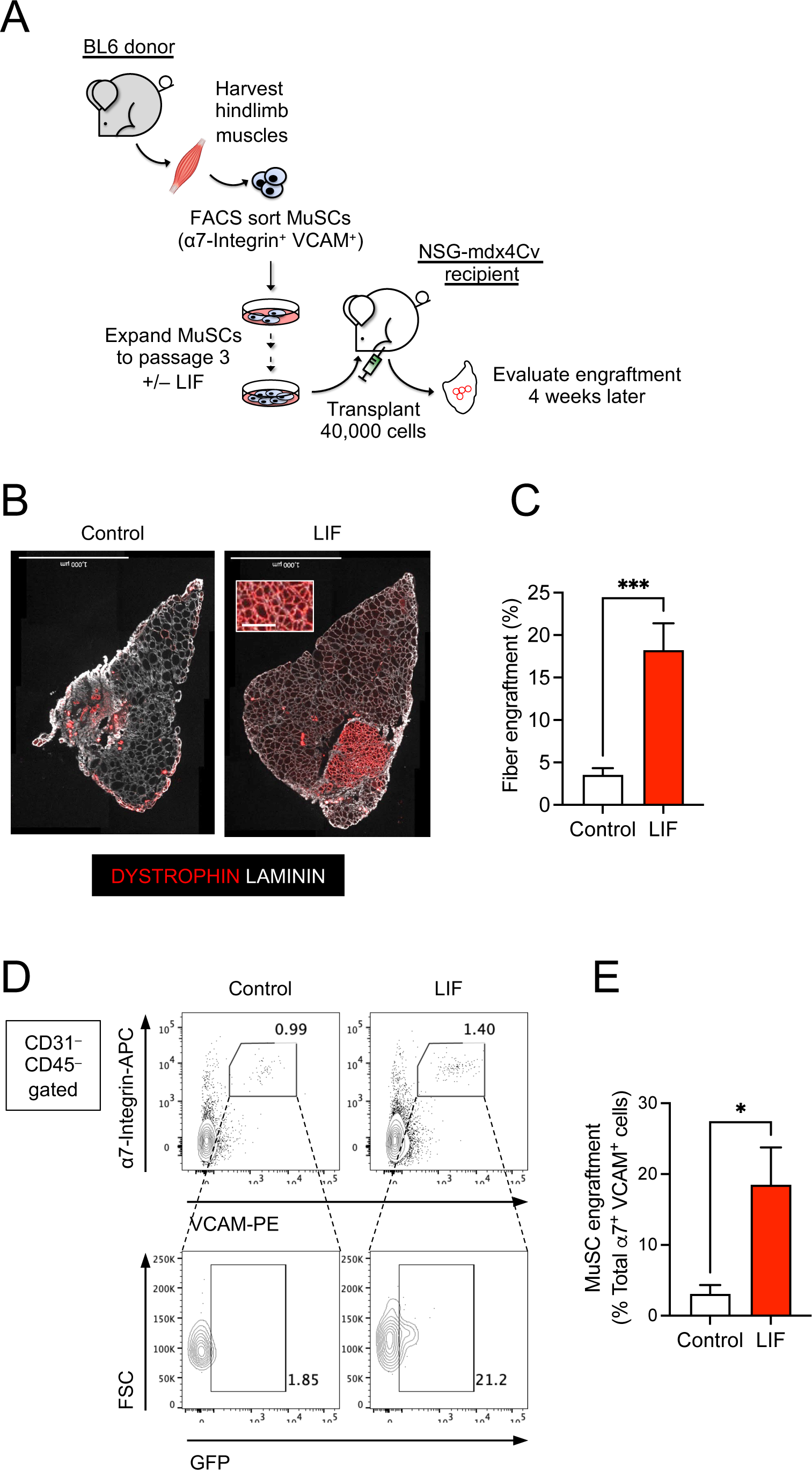
LIF enhanced the potential of cultured MuSCs to develop into muscle fibers and repopulate the endogenous MuSC niche upon transplantation. (**A**) Schematic of the evaluation of the effect of LIF treatment on the engraftability of cultured MuSCs. (**B**) LIF-treated passage 3 MuSCs engrafted and formed new DYSTROPHIN^+^ fibers. Total fibers (donor-derived and recipient) are indicated as LAMININ^+^ (scale bar = 1000 μm). (**C**) Quantification of (**B**) (mean ± SEM, 8 biological replicates, *** p < 0.001). (D) FACS analysis showing LIF-treated passage 3 MuSCs engrafted and repopulated the MuSC compartment. (**E**) Quantification of (**D**) (mean ± SEM, 3 biological replicates, * p < 0.05).

We next evaluated whether LIF-treated MuSC cultures could repopulate the endogenous MuSC pool. For this experiment, we used H2B-GFP BL6 mice as donors to obtain α7-Integrin^+^ VCAM^+^ MuSCs. These mice constitutively express GFP in their nuclei and thereby allow distinction between donor (GFP^+^) and recipient (GFP^−^) cells upon transplantation. FACS-sorted GFP^+^ MuSCs were subsequently cultured with or without LIF (1000 units/mL) for 12 days over 3 passages for transplantation. Four weeks after transplantation, we observed a significant contribution of donor-derived cells in the MuSC compartment (GFP^+^ α7-Integrin^+^ VCAM^+^) (Figure 3D-E). Therefore, these results confirmed the role of LIF-LIFR signaling in regulating the engraftment of cultured MuSCs to develop into muscle fibers and repopulate the endogenous MuSC niche.

## 4. Discussion

A major obstacle in developing cell therapy to treat muscular dystrophies is poor engraftment outcome of donor cells. Whereas endogenous muscle stem cells have tremendous regenerative potency when they are transplanted right away after isolation, their engraftability abruptly diminish once they are expanded *in vitro* (Montarras et al., 2005; Sacco et al., 2008; Xie et al., 2021). Despite advancements such as p38 modulation and extracellular matrix modification, muscle stem cells are unable to engraft robustly beyond 7 days in culture (Charville et al., 2015; Gilbert et al., 2010; Parker et al., 2012; Quarta et al., 2016). We need a new strategy to discover factors that can improve the engraftment potential of cultured muscle stem cells.

We recently demonstrated that skeletal myogenic progenitors from PSC *in vivo* differentiation had remarkable regenerative potency, and their engraftment potential was on par with bona fide endogenous muscle stem cells (Chan et al., 2018; Xie et al., 2023). Importantly, these skeletal myogenic progenitors remain engraftable even after prolonged *in vitro* culture (Xie et al., 2021). The superior engraftability and expandability of *in vivo* differentiated skeletal myogenic progenitors make them a novel platform for identifying factors that regulate engraftment. In this regard, we have identified LIF-LIFR signaling as a potential engraftment-regulatory pathway. The expression of LIFR was positively correlated to engraftability: highest expression in the highly engraftable cells, modest expression in the moderately engraftable cells, and minimal expression in the less engraftable cells. We subsequently showed that LIF, an endogenous ligand for LIFR, upregulated the muscle stem cell factor PAX7 in cultured muscle stem cells and increased the number of PAX7^+^ “reserve cells” in differentiated cultures. More importantly, LIF improved the engraftment potential of muscle stem cells that had been cultured and expanded *in vitro* for 12 days by 5-fold. This is particularly interesting because prior research primarily examined the role of LIF signaling in regulating MuSC functions during highly regenerative conditions such as injured muscles *in vivo*, i.e., LIF activation further accelerated the already proficient regenerative capacity of endogenous MuSCs (Austin et al., 2000; Kami et al., 2000; Kami and Senba, 1998; Tham et al., 1997; White et al., 2001). In contrast, our current study focused on cultured/expanded MuSCs that have already lost their engraftment potential. Our work thus supplemented previous findings.

Despite numerous efforts, the molecular mechanisms that regulate whether a given transplanted muscle cell population is engraftable remains unclear. In the current study, we have demonstrated the feasibility of our *in vivo* differentiation platform for identifying novel factors that regulate engraftability. Specifically, we have successfully used our platform to discover the beneficial role of LIF-LIFR signaling in improving the ability of cultured muscle stem cells to form new fibers in transplantation assays. Future investigations might reveal additional candidates for further promoting the engraftability of donor muscle cells.

## Author Contributions

Conceptualization: S.S.K.C.; Methodology: N.X. and S.S.K.C.; Investigation: N.X., K.R., T.S. and S.S.K.C.; Formal Analysis: N.X., K.R. and S.S.K.C.; Visualization: N.X., K.R. and S.S.K.C.; Writing: N.X., K.R. and S.S.K.C.; Funding Acquisition: S.S.K.C.; Supervision: S.S.K.C.

## Acknowledgments

The monoclonal antibody to PAX7, embryonic MHC, neonatal MHC, MHC, MHC-I and MHC-IIa were obtained from the Developmental Studies Hybridoma Bank developed under the auspices of the NICHD and maintained by the University of Iowa.

## Data Availability Statement

Publicly available datasets were analyzed in this study. This data can be found here: GEO: GSE182508.

